# Nectar- and stigma-specific expression of a chitinase could partially protect against fire blight in certain apples

**DOI:** 10.1101/606046

**Authors:** Anita Kurilla, Timea Toth, Laszlo Dorgai, Zsuzsanna Darula, Tamas Lakatos, Daniel Silhavy, Zoltan Kerenyi, Geza Dallmann

**Affiliations:** Agricultural Biotechnology Institute, Szent-Györgyi 4, H-2100, Gödöllo, Hungary; Research Institute for Fruitgrowing and Ornamentals, Park 2, H-1223, Budapest, Hungary; Biocenter Ltd, Temesvári 62, H-6726, Szeged, Hungary; Hungarian Academy of Sciences, Biological Research Centre, Temesvári krt 62, H-6726 Szeged, Hungary; MTKI, Lucsony 24, H-9200, Mosonmagyaróvár, Hungary

**Keywords:** acidic chitinase, antibacterial effect, *Erwinia amylovora*, fire blight tolerance, *Malus floribunda* 821, MYB305, nectar-and stigma-specific expression, promoter repeat

## Abstract

To attract pollinators many angiosperms secrete stigma exudate and nectar in their flowers. As these nutritious fluids are ideal infection points for pathogens, both secretions contain various antimicrobial compounds. *Erwinia amylovora*, the causing bacterium of the devastating fire blight apple disease, is the model pathogen that multiplies in flower secretions and infects through the nectaries. Although *Erwinia* resistant apples are not available, certain cultivars are tolerant. It was reported that in stigma infection assay, the ‘Freedom’ cultivar was *Erwinia* tolerant while the ‘Jonagold’ was susceptible. We hypothesized that differences in the nectar protein compositions lead to different susceptibility. Indeed we found that an acidic chitinase III protein (Machi3-1) selectively accumulates in the nectar and stigma of the ‘Freedom’ cultivar. We demonstrate that MYB binding site containing repeats of the ‘Freedom’ *Machi3-1* promoter are responsible for the strong nectar- and stigma-specific expression. As we found that *in vitro* the Machi3-1 protein impairs growth and biofilm formation of *Erwinia* at physiological concentration, we propose that the Machi3-1 contribute to the tolerance by inhibiting *Erwinia* multiplication in the stigma exudate and in the nectar. We show that the *Machi3-1* allele was introgressed from *Malus floribunda* 821 into different apple cultivars including the ‘Freedom’.

**Highlight:** Certain apple cultivars accumulate to high levels in their nectar and stigma an acidic chitinase III protein that can protect against pathogens including fire blight disease causing *Erwinia amylovora*

## Introduction

Plants secrete rewarding, sugar rich fluids such as stigma exudates and nectar to attract pollinators (Tanveer *et al*., 2014). Based on their stigma Angiosperm can be divided into two groups, plants with dry (E.g.: *Arabidopsis thaliana*, Arabidopsis) and wet stigma (E.g.: *Malus domestica*, apple and *Nicotiana tabacum*, tobacco). While the surface of the dry stigma is covered with a proteinaceous extracuticular layer (pellicle), the wet stigma is covered with stigma exudates (Edlund *et al*., 2004). This fluid secretion plays a critical role in pollen capture, hydration, growth of pollen tube and serves as a reward for the pollinators. The exudate can also be found at the intercellular spaces of the stigmatic zone and transmitting tissue in mature pistils (Rejón *et al*., 2014). Floral nectars are secreted by specific glands called nectaries (Heil, 2011; Roy *et al*., 2017). Nectaries have evolved independently multiple times, and they can differ in their position, morphology, and secretion mechanism (De la Barrera and Nobel, 2004). Despite these differences, nectary development is conserved in most angiosperms (Min *et al*., 2019). The C-lineage genes regulate the expression of the Crab Claws transcription factor, which is essential for nectary development (Bowman and Smyth, 1999; Lee *et al*., 2005; Morel *et al*., 2018). Jasmonic acid (JA) is required for nectar secretion, while auxin mainly controls the volume of the nectar (Radhika *et al*., 2010; Bender *et al*., 2013; Roy *et al*., 2017). JA induces nectary-specific gene expression by stimulating the degradation of JAZ proteins (Kelley and Estelle, 2012), thereby releasing the JAZ repressed transcriptional factor MYB305 or the homologs of MYB305 (Liu *et al*., 2009; Liu and Thornburg, 2012; Stitz *et al*., 2014; Schmitt *et al*., 2018*a*). MYB305 directly and indirectly promotes the transcription of nectary-specific genes, including nectarins, genes whose proteins products are secreted into the nectar (Liu and Thornburg, 2012). The chemical composition of the stigma exudate and the nectar is relatively different but both secretions are highly nutritious containing high levels of sugars. The stigma exudate is rich in complex sugars and proteins and contains at lower concentration free sugars, amino acids and lipids (Pusey *et al*., 2008). By contrast, the nectar is rich in sucrose and hexoses and free amino acids. It also contains additional components such as phenolics, secondary metabolites and nectarins (Heil, 2011). The proteome of the floral nectars is relatively simple and frequently contains only a few (sometimes only one or two) dominant proteins (Roy *et al*., 2017). The protein profile of the stigma exudates is more complex suggesting that it is a physiologically more active extracellular fluid (Rejón *et al*., 2013). Both secretions are excellent medium for microbes. Microbial infection of the flower secretions is harmful as microbes can alter the chemical composition of the fluids, and mainly because plant pathogens can infect efficiently by multiplying in these nutritious fluids and then by entering into the plant through the stomata of the nectaries. Thus it is not surprising that plants accumulate antimicrobial components including antimicrobial proteins (such as chitinases and glucanases) in these secretions. Indeed, in the proteome of stigma exudates, the defense and stress response proteins are the dominant GO categories (Sang *et al*., 2012). Flower nectars frequently contain antimicrobial proteins in very high concentrations (Zha *et al*., 2016; Ma *et al*., 2017; Nogueira *et al*., 2018; Schmitt *et al*., 2018*b*) or accumulate nectarins that generate antimicrobial hydrogen peroxide in the nectar (Carter *et al*., 2007). Furthermore, it was shown that nectar of wild squash is antibiotic and efficiently reduces the symptoms of the bacterial wilt (Sasu *et al*., 2010). The gram-negative bacterium *Erwinia amylovora* that is one of the most devastating bacterial pathogens of apple, is the classical example of pathogens that multiply in flower secretions and infects through the nectaries (Farkas *et al*., 2012; Malnoy *et al*., 2012). *Erwinia* first colonizes the stigma and multiplies in the stigma exudates, then the pathogen is washed down by rain or dew into the nectar. The pathogen further multiplies in the nectar and finally enters into the plant through the stomata of the nectaries (Bubán *et al*., 2003). *E*. *amylovora* produces exopolysaccharides that are involved in biofilm formation (Koczan *et al*., 2009). Pear fruit and apple shoot inoculation assays show that mature biofilm formation is needed for full virulence of *Erwinia* (Koczan *et al*., 2011; Piqué *et al*., 2015). Although *Erwinia* resistant apple cultivars are not available, certain cultivars are tolerant (Gusberti *et al*., 2015). These cultivars are infected less frequently and develop reduced symptoms. For instance, after inoculation of the stigmas of the tolerant ‘Freedom’ and the susceptible ‘Jonagold’ cultivars, ‘Freedom’ was less infected, much less bacteria were detected on the surface of the nectaries and tissue coloring symptoms were much weaker on the ‘Freedom’ (Mihalik *et al*., 2004). It was assumed that the chemical composition of the ‘Freedom’ and ‘Jonagold’ nectars was identical, and proposed that the rough surface of the ‘Jonagold’ nectary was responsible for the more efficient colonization and the stronger symptoms (Mihalik *et al*., 2004). However, accumulating data indicate that nectarins can play important antimicrobial role (Heil, 2011; Roy *et al*., 2017). Therefore we wanted to test an alternative (but not mutually exclusive) hypothesis that the nectar protein profiles of the tolerant and susceptible cultivars are different. Indeed, we found that an acidic chitinase III protein (Machi3-1) accumulates to high level in the nectar and the stigma of the tolerant ‘Freedom’ cultivar but not in the susceptible cultivars. We show that different *Machi3-1* alleles are present in ‘Freedom’ and ‘Jonagold’ cultivars and that the presence of 5 direct repeats in the promoter of ‘Freedom’ *Machi3-1* allele is responsible for the strong nectar- and stigma-specific expression. We demonstrate that the strongly expressing *Machi3-1* allele was introgressed from *Malus floribunda* 821 into different cultivars including ‘Freedom’. Relevantly, we found that Machi3-1 protein can inhibit the growth and biofilm formation of *E*. *amylovora in vitro* at physiological concentration. How the stigma- and nectar-specific expression of *Machi3-1* could contribute to the *Erwinia* tolerance and in general to plant defense will be discussed.

## Materials and methods

### Bacterial strain and plant materials

The bacterial strain *Erwinia amylovora* ref T was grown in TSB (Tryptic Soy Broth) medium at 28°C overnight. The plant materials were collected from the cultivar collection of the Research Institute for Fruitgrowing and Ornamentals (Újfehértó, Hungary) from 2005 to 2015. We used various scab resistant and susceptible cultivars. *Malus domestica* Borkh. cultivars ‘Jonagold’, ‘Sampion’, ‘Golden Delicious’, ‘Gala’ ‘Idared’, ‘Redwinter’, ‘Red Rome’ and the Hungarian landrace ‘Simonffy’ are scab susceptible, while Releika, Resi, Remo, Rewena (Germany) ‘Rajka’, ‘Selena’, ‘Topaz’, ‘Rubin’, ‘Rubinola’ (Czech Republic), ‘Hesztia’ (Hungary) are scab resistant cultivars. F1 hybrids of ‘Freedom’ x ‘Redwinter’ and ‘Freedom’ x ‘Red Rome’ derived from earlier crossbreeding program.

### Nectar and stigma collection

Apple nectars were collected from field grown plants. Nectars from transgenic tobaccos, which were grown in the greenhouse, were collected in the morning (9-10 am). Nectars from the same cultivar were usually pooled and stored at −70°C. Stigma samples were pooled from 5 stigmas of the same plant.

### Nucleic acid techniques and protein extraction

Genomic DNA was purified with Quick-DNA Plant/Seed Miniprep kit (Zymo Research D6020). RNA extraction was carried out as described (Szittya *et al*., 2002). To purify plant protein extract, 100 mg plant tissue was homogenized with 400 µl extraction buffer (100 mM NaCl, 100 mM glycine, 10 mM EDTA, 2 % SDS), incubate at 95°C for 5 minutes and centrifuged. Protein concentration was measured at 280 nm.

### Stain-free protein profiles and western-blot assays

Nectars and the protein extracts were separated by stain-free 1D SDS-PAGE (Bio-Rad’s Mini PROTEAN® TGX Stain-Free™ Gels). For western blot assays, samples were separated by SDS-PAGE, blotted onto Amersham Protran membrane (GE Healthcare, 10600008) and hybridized with rabbit polyclonal antibody serum raised against Machi3-1. ECL Anti-Rabbit IgG Horseradish Peroxidase linked (GE Healthcare, NA934-1ML) secondary antibody was used for detection. Actin antibody (Anti-Actin Plant MerckA0480) was used for control. Chemiluminescent protein detections were conducted with ECL Western Blotting Substrate (Promega, W1001), according to the manufacturer’s instructions. Western blots were scanned with ChemiDoc MP System and analyzed with ImageLab 5.0 software (Bio-Rad).

### Protein sequencing

The dominant protein band of ‘Freedom’ nectar was partially sequenced (described in details at Supplementary Materials and Methods S1). Briefly, the excised protein band was in-gel digested as described (Migh *et al*., 2018). Peptides were analyzed by data-dependent LC-MS using a Waters Q-TOF Premier mass spectrometer online coupled to a nanoAcquity uHPLC system. Raw data was converted into a peaklist using the ProteinLynx PLGS software and the data was searched using the Batchtag Web software of the Protein Prospector search engine. As automated protein identification did not yield high confidence identifications, MS/MS data were inspected manually and high-quality MS/MS spectra were evaluated manually. Protein segments were used for degenerate PCR primer designing.

### PCR cloning of the Machi3-1 gene from Freedom cultivar

Degenerated oligonucleotides (*aldegf* and *aldegr*, respectively) were designed for the predicted N-proximal ADYIWNNF and the C-proximal WNRFYDN peptide segments. cDNA was prepared from total RNA isolated from the nectary rich tissues of ‘Freedom’. PCR product was amplified, subsequently cloned and sequenced. Based on this information specific oligonucleotides were synthesized to clone the genomic region by inverse PCR (*invj1 for, invj2 for, invba1 rev, invba2 rev*).

### RT-PCR assays

For quantitative RT-PCR, total RNAs were treated with DNase I (Thermo Fisher Scientific, EN0525), and cDNAs were transcribed using a RevertAid First Strand cDNA Synthesis kit (Thermo Fisher Scientific, K1621). qRT-PCR was carried out with Fast Start Essential DNA Green Master Mix (Roche, 06402712001) in a Light Cycler 96 Real-Time PCR instrument (Roche). For semi-quantitative RT-PCR assays, the same cDNAs were used in conventional PCR reactions using DreamTaq Green PCR Mastermix (Thermo Fisher Scientific, K1081) in a T-Personal thermal cycler (Biometra).

### Machi3-1 genotyping

The promoter regions of *Machi3-1* alleles were amplified with the *SaFreeFor* and *SaFreeRev* primers and separated on 1.5 % agarose gel.

### Cloning of the transgenic constructs

To generate 5B-Machi3-1 and 2B-Machi3-1 transgenic constructs, the promoter and coding region of *5B-Machi3-1* and *2B-Machi3-1* genes were amplified with the *5B2BproFor* and *Machi3-1-stopRev* primers, and then the PCR products were cloned into *HindIII* and *BamHI* cleaved Bin61S vector (Silhavy *et al*., 2002). To create promoter deletion constructs, promoter segments were amplified using one of the *1*.*2kbFor, 1kbFor, 0*.*9kbFor, 0*.*6kbFor, 0*.*4kbFor, 0*.*2kbFor* forward primers and the *Machi3-1stopRev* reverse primer. The fragments were cloned into *HindIII* and *HpaI* cleaved 5B-Machi3-1 transformation vector to replace the original promoter regions. The constructs were sequenced. The list of primers are shown as Supplementary Table S1.

### Plant transformation

Leaf disc transformation was carried out to generate transgenic *N.tabaccum* plants (Bevan *et al*., 1985). Transgenic tobaccos were selected on kanamycin containing media and then the regenerants (T_0_ plants) were grown in the greenhouse.

### Expression of recombinant proteins in Pichia pastoris, analysis of protein expression

Machi3-1 protein was expressed with Pichia Expression Kit (Invitrogen K1710-01) using pPICz vector. The protein was expressed according to the manufacturer’s protocol. The signal peptide of Machi3-1 transports the protein to the extracellular space, allowing the purification of the protein from the supernatant. The secreted protein was purified from the supernatant by precipitating with ammonium-sulfate (60% saturation). The precipitate was pelleted (15000 rpm, 10 min., 4°C), resuspended in 5 ml 10 mM Sodium-acetate (pH 5.0) and concentrated with Amicon Ultra-4 Centrifugal Filter Units (Merck Millipore, 10K) ∼10 fold, reaching the final protein concentration ∼3 mg/ml. Empty pPICz vector transformant *P.pastoris* was grown and induced as the test strain, and then its supernatant was similarly treated (ammonium-sulfate precipitated, centrifuged, resuspended in 10 mM Sodium-acetate and concentrated to ∼3 mg/ml). The purified supernatant of the empty vector transformant *P.pastoris* was used as negative control in activity assays. SDS-PAGE assay was used to test that the background of the negative control and the purified Machi3-1 was similar (Fig. S2A and S2B).

### Chitinase and lysozyme activity assay

Chitinase activity was measured by Schales’ reagent method (Ferrari *et al*., 2014). Colloid chitin was prepared according to Shen (Shen *et al*., 2010) with minor modifications. 6 g chitin was suspended in 200 ml 37 % HCl and agitated overnight at 4°C. 1 L of distilled water was added followed by centrifugation at 8000 g for 20 minutes. The pellet was washed with water till the pH reached 5.0.

Colloid chitin (at 3 mg/ml final concentration) was incubated in 200 µl of 50 mM KPO_4_ (pH 6.0.) with increasing amounts (50-400 ng) of Machi3-1 protein. The reactions were rotated at 30°C for 1 hour at 200 RPM. Samples were briefly centrifuged (10 sec) and 100 µl supernatant was transferred to a new tube. 100 µl Schales’ reagent (0.5 M sodium carbonate and 0.5 g/L potassium ferricyanide in water) was added and then boiled at 97°C for 15 minutes. After cooling down to RT, absorbance was measured at 420 nm. Chitinase from *Streptomyces griseus* (Sigma, 9001-06-3) was used for positive control. Purified supernatant of the empty vector transformant *P.pastoris* was used as a negative control.

Lysozyme activity was measured by agar diffusion plate method. *Micrococcus lysodeikticus* (Merck, 4698) was used as the substrate (0.05 mg/ml). 1 % agarose gel containing 1 mg *M.lysodeikticus in* 10 mM sodium-acetate buffer (pH 5.0.) was made (20 ml per plate). After solidification, lysozyme (Merck L6876) or Machi3-1 were loaded into the wells (4 mm diameter). Plates were incubated at 30°C for 24 hours.

### In vitro Erwinia growth inhibition assay

Bacterial *in vitro* growth inhibition assay was carried out with minor modificiations as described (Nash *et al*., 2006). Approximately 10^2^ *E.amylovora* cells were suspended in 200 µl of 10 mM PBS (pH 7.4). The suspension was incubated without shaking for 24 hours at 28°C with different amount of purified Machi3-1 protein, or with purified supernatant of empty vector transformant *P.pastoris* as a negative control. Viable cells were counted by plating.

### Enzymatic detachment of Erwinia biofilm

*In vitro* biofilm detachment assay was monitored with crystal violet staining (Koczan *et al*., 2009; O’Toole, 2011). *E.amylovora* overnight culture was diluted in LB to 1:100 and 130 µl of the diluted culture was incubated at 30°C in a 96-well TC-treated Tissue culture polystyrene plate (1 × 10^6^ cells per well) to allow biofilm formation. After 24 hours the suspension was removed, and then 130 µl of purified Machi3-1 diluted in 50mM KPO_4_ buffer (pH 6.0) was added to the biofilm covered plates. The reactions were kept at 28°C for 3 hours. Wells were washed 3 times with dH_2_O, then 150 µl 0.1% crystal violet (CV) was added. After 15 minutes CV was removed, then the plate was washed 3 times with dH_2_O, and dried for overnight. For quantification 30% acetic acid was added to each well, incubated for 15 minutes and the OD was measured at 550 nm.

### Statistics

Bacterial growth inhibition assays were repeated four times in independent experiments. Biofilm detachment experiments were performed in octuplicate and repeated three times. Comparisons between groups were done by ANOVA and Tukey-test to determine P-values. Statistical significance * was set at p<0,05 and *** p<0,001.

### Bioinformatical analysis

Sequence analysis was made by BLASTN and BLASTP softwares. Transcription factor binding sites were predicted using PlantTFDB (Plant Transcription Factor Database) (Jin *et al*., 2017). Protein and DNA sequences were aligned by ClustalW method using the MegAlign program. Structural alignment and homology modelling of Hevamine as the template and the Machi3-1 protein was carried out by the SPDBV (Swiss-PdbViewer) program. Phylogeny tests were made using Bootstrap method (No. of Bootstrap Replications = 1000) and analyzed by UPGMA statistical method using the MEGA6 software.

## Results

### A class III chitinase-like protein accumulates in the nectar of the Erwinia tolerant ‘Freedom’ apple cultivar

To test our hypothesis that the nectar composition of the fire blight tolerant and susceptible apple cultivars is different, nectar protein profiles of the tolerant ‘Freedom’, the susceptible ‘Jonagold’ and ‘Sampion’ cultivars were studied by 1D SDS-PAGE. None of the nectar proteins accumulated to high levels in the susceptible cultivars, while the nectar of ‘Freedom’ contained a 29kDa dominant protein (Fig. 1A). Although this protein was present at very high concentration (∼50-80 ng/µl) in the ‘Freedom’ nectar, it was not detectable in the nectars of the susceptible cultivars (Fig. 1A). The analysis was repeated in four consecutive years with the same results, therefore the presence of this dominant protein in the ‘Freedom’ nectar was not due to any environmental condition. The 29kDa protein was isolated, partially sequenced, then primers were designed and inverse PCRs were conducted to clone the genomic copy of the gene from the ‘Freedom’ cultivar. The amplified region contained a 894 nucleotide (nt) long intronless coding sequence, a long (1417 nt) upstream and a short (77 nt) downstream regions. Sequence analysis revealed that the predicted ‘Freedom’ nectar protein is a class III chitinase (will be referred to as Machi3-1 for *<u>Ma</u>lus* <u>chit</u>inase<u>III</u>-<u>1</u>). Class III chitinases belong to the GH18 endochitinase family (Adrangi and Faramarzi, 2013). Machi3-1 is an acidic class III chitinase (calculated isoelectric point is 4.4), which shows strong sequence similarity (66.4%) to the well characterized class III chitinases as PSC (pomegranate seed chitinase) and Hevamine (64.18%) (Terwisscha Van Scheltinga *et al*., 1996; Lv *et al*., 2011; Masuda *et al*., 2015). The critical catalytic amino acids and the cis-peptides (involved in chitin binding) are all conserved. Moreover, homology modelling predicts that the structure of Machi3-1 protein is highly similar to the structure of Hevamine (Fig. S1). Machi3-1 contains an N-terminal signal peptide that destines proteins towards the secretory pathways (Chung and Zeng, 2017). These data suggest that the Machi3-1 protein is a functional, secretable acidic chitinase.

**Fig 1.**
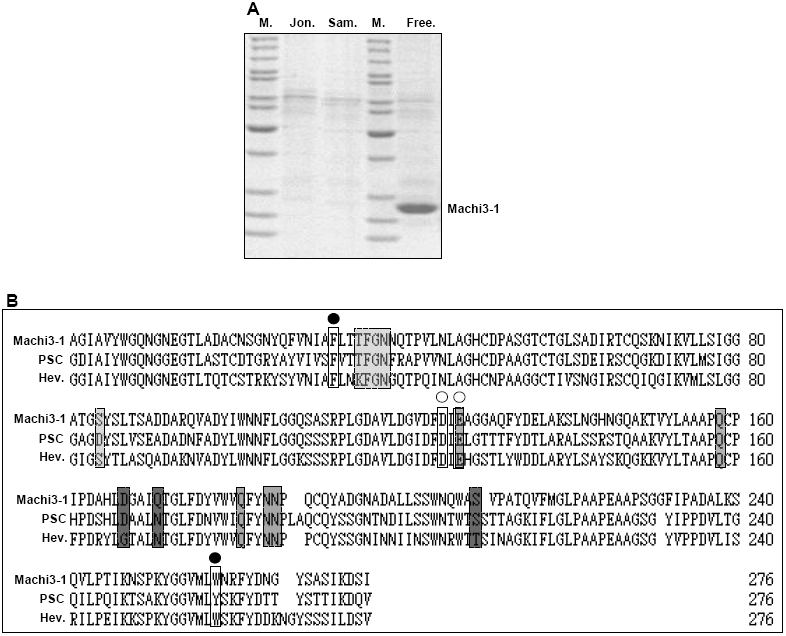
Machi3-1 acidic chitinase III protein accumulates to high levels in the nectar of ‘Freedom‘apple cultivar. (A) The nectar protein profile of ‘Jonagold’ (Jon.), ‘Sampion’ (Sam.) and ‘Freedom’ (Free.) cultivars were studied by SDS-PAGE. Note that Machi3-1 protein accumulates only in the ‘Freedom’ nectar. M. shows size marker. (B) Multiple sequence alignment of Machi3-1 with PSC (Pomegranate seed chitinase) and Hevamine. The N-terminal signal peptide regions were omitted from the alignment. Substrate binding cleft are shown as •, while the active site is marked with ○. The three magnesium binding sites of PSC (3 to 5 amino acid/binding site) are shown by empty, light green and dark grey columns.

### Machi3-1 is an active chitinase

Basic class III chitinases frequently have dual chitinase and lysozyme activities, while the acidic class III chitinases have strong chitinase but only weak or no lysozyme activity (Ma *et al*., 2017). To characterize the Machi3-1 protein, it was expressed in *P.pastoris* and the chitinase and lysozyme activities of the purified protein were tested *in vitro*.

To measure the chitinase activity of the purified Machi3-1 protein, Schales’ procedure using colloidal chitin for a substrate was carried out (Ferrari *et al*., 2014). *S.griseus* chitinase and the supernatant of empty vector transformed *P.pastoris* were used as positive and negative controls, respectively. Machi3-1 proved to be a relatively efficient chitinase; its activity was ∼25% of the *S.griseus* chitinase (Fig. 2A). The Machi3-1 had a barely detectable lysozyme activity in *Micrococcus* lysis assays (Fig. S2C). Thus we concluded that Machi3-1, like most acidic chitinase III proteins, has strong chitinase and very weak lysozyme activity.

**Fig 2.**
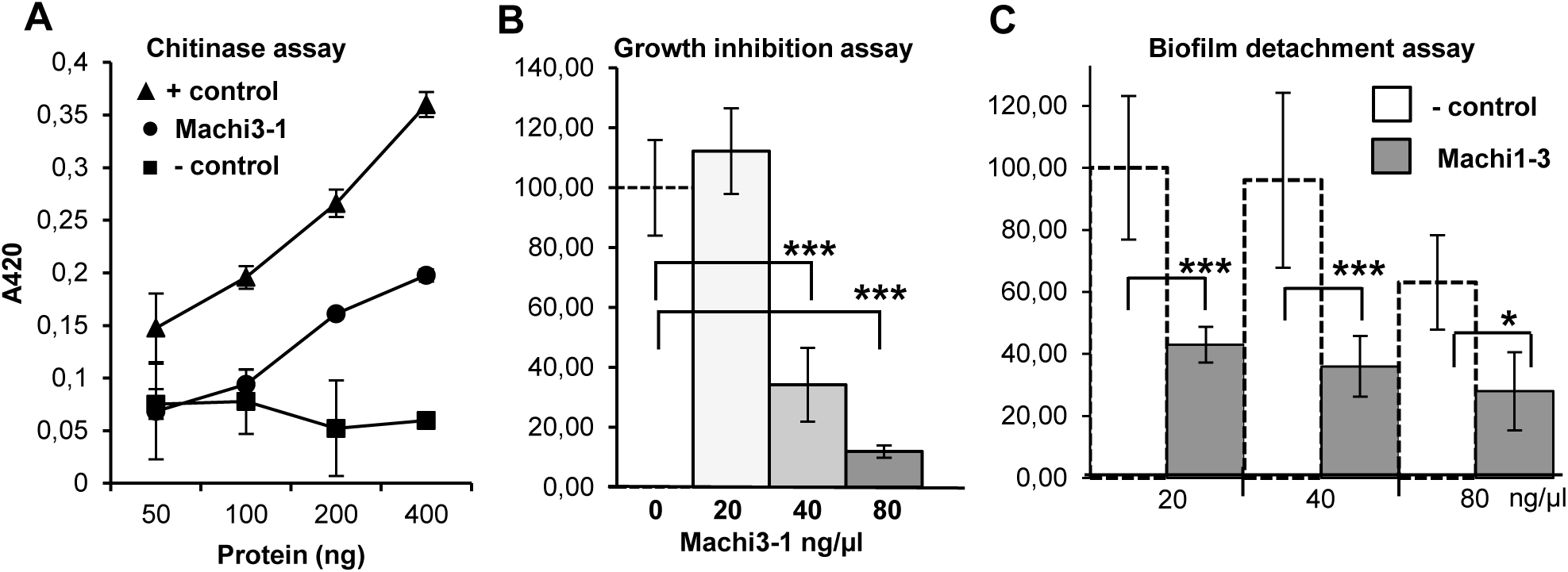
Machi3-1 has an antibacterial effect against *E.amylovora*. (A) *In vitro* chitinase assay was conducted with purified Machi3-1 protein and with *S.griseus* chitinase as positive and with supernatant from empty vector transformed strain as negative control. Note that the positive control is ∼ 4 times more effective than the Machi3-1. (B) Machi3-1 inhibits the growth of *E.amylovora*. (C) Biofilm detachment assay shows that Machi3-1 impairs biofilm formation. Significancy levels: * *p*-value <0.05, *** *p*-value <0.001.

### Machi3-1 inhibits growth and biofilm formation of E.amylovora in vitro

Next we tested if Machi3-1 had an antibacterial effect against *E.amylovora*. Bacterial cultures were incubated with Machi3-1 protein purified from *P.pastoris* supernatant or with supernatant of empty vector transformant *P.pastoris* for a negative control. We found that at high concentration (40-80 ng/µl) Machi3-1 significantly reduced the growth of *E.amylovora* (Fig. 2B). Relevantly, the Machi3-1 protein is present in the nectar of Freedom cultivar in similar ∼50-80 ng/µl concentration (Fig. S2B).

Biofilm formation is required for successful *Erwinia* infection (Koczan *et al*., 2011). As chitinases can impair biofilm formation (Chung *et al*., 2014), we wanted to study if Machi3-1 modifies biofilm formation of *E.amylovora*. Pre-formed *E.amylovora* biofilm was treated with purified Machi3-1 protein, and then biofilm detachment was quantified by crystal violet staining assay (O’Toole, 2011). As Fig. 2C shows, the Machi3-1 efficiently detached the preformed *E.amylovora* biofilm at physiological concentration. Taken together, the Machi3-1 acidic chitinase III protein efficiently impairs the growth and biofilm formation of *E.amylovora in vitro* at physiological concentration.

### Expression of Machi3-1 gene in ‘Freedom’ and ‘Jonagold’ cultivars

To analyze the expression pattern of *Machi3-1* gene, polyclonal antibody was produced, and accumulation of the Machi3-1 protein was studied in different tissues of the ‘Freedom’ and ‘Jonagold’ cultivars (Fig. 3A and Fig. S3). Confirming our earlier data, the Machi3-1 protein accumulated to high levels in the Freedom nectar but was barely detectable in the Jonagold nectar (Fig. S3). Moreover, in the ‘Freedom’ cultivar the Machi3-1 protein was also abundant in the stigma tissue but accumulated to low levels in the nectary, leaf, petal, stamen and ovary samples. In the ‘Jonagold’ cultivar, the Machi3-1 protein accumulated to low levels in all samples (Fig. 3A). Next we studied the expression of *Machi3-1* at mRNA level in the nectary, stigma and leaf samples of the two cultivars (Fig. 3B). In the ‘Freedom’ cultivar, *Machi3-1* transcript expressed to very high levels in both the nectary and the stigma but it was barely detectable in the leaves. By contrast, *Machi3-1* mRNA accumulated to low levels in all ‘Jonagold’ samples (Fig. 3B). Taken into consideration that (i) *5B-Machi3-1* mRNA expressed in the ‘Freedom’ nectary, while the protein accumulated in the nectar, and that (ii) the Machi3-1 protein has an export signal, we conclude that the *5B-Machi3-1* is a nectarin gene, which expresses in the nectary cells and then its protein product is secreted into the nectar.

**Fig 3.**
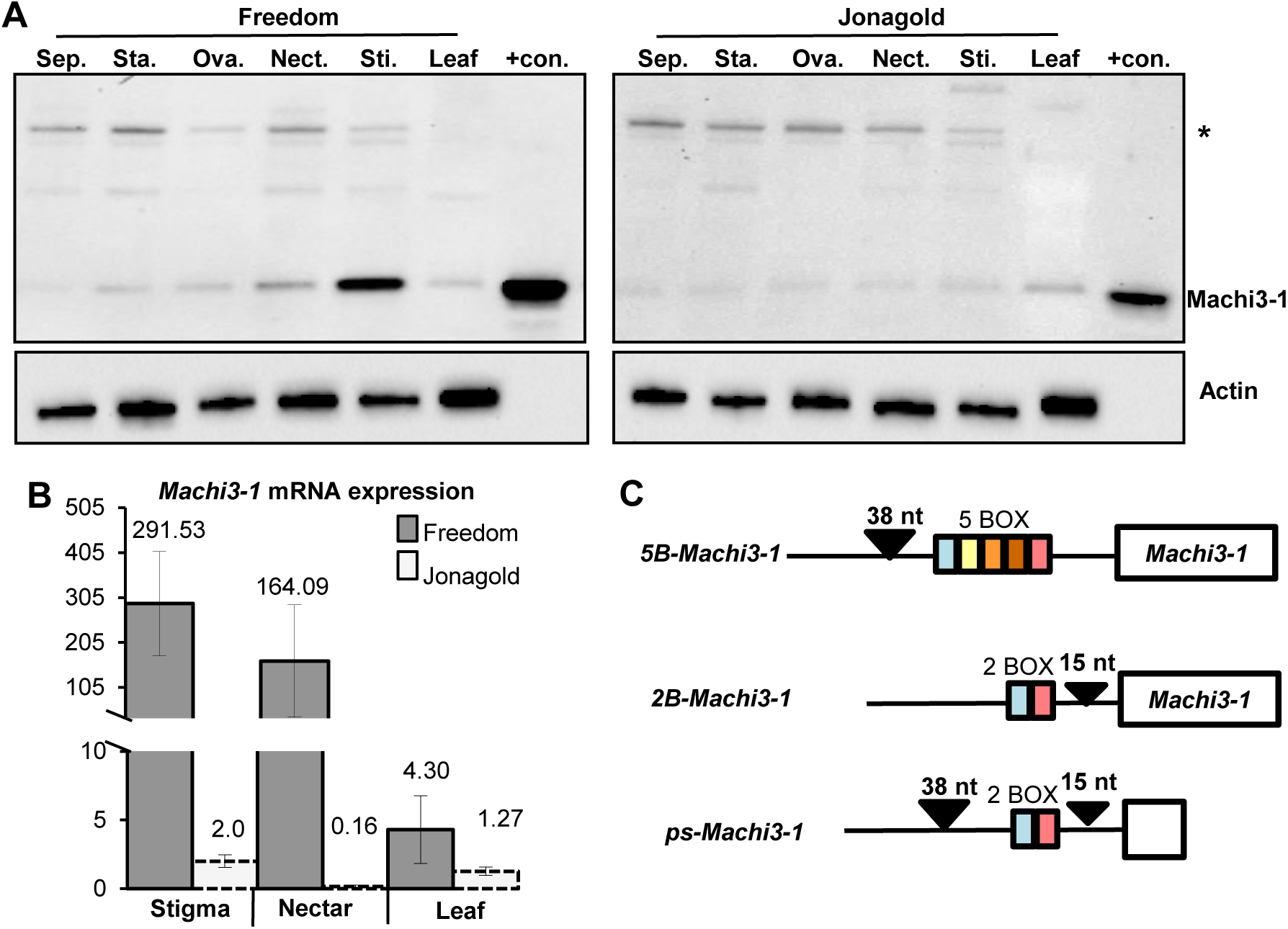
Expression of Machi3-1 in ‘Freedom’ and ‘Jonagold’ apple cultivars. (A) Machi3-1 protein is abundant in the stigma of ‘Freedom’ cultivar. Western-blot assay was conducted to monitor the expression of Machi3-1 protein in leaf, stigma (Sti.), nectary (Nect.), ovary (Ova.), stamen (Sta.) and sepal (Sep.) samples in ‘Freedom’ and ‘Jonagold’ cultivars. Actin probe was used as loading control. ‘Freedom’ nectar was used as positive control (+con.) for the Machi3-1 blot. Note that Machi3-1 accumulates to low levels in the ‘Freedom’ nectary but it is very abundant in the nectar and that actin, which lacks signal peptide, does not accumulate in the nectar. (B) Expression of *Machi3-1* mRNA. Quantitative RT-PCR assay was conducted to study the expression *Machi3-1* mRNA in different organs of ‘Freedom’ and ‘Jonagold’ cultivars. (C) Non-proportional schematic representation of the different *Machi3-1* alleles. White box shows the coding region. Black triangles indicate the allele specific insertions. The differently colored boxes represent the direct repeats. Note that *ps-Machi3-1* is a pseudogene.

### The promoter regions of the ‘Freedom’ and ‘Jonagold’ Machi3-1 alleles are different

We postulated that variations in the *Machi3-1* promoters are responsible for the strikingly different mRNA expression between ‘Freedom’ and ‘Jonagold’ cultivars. Therefore, the coding and the promoter regions of the *Machi3-1* gene were also cloned from the ‘Jonagold’ cultivar, and then the ‘Freedom’ and ‘Jonagold’ *Machi3-1* genes were compared. The nucleotide sequences of the coding regions and the predicted protein sequences are almost identical (only 4/894 nt and 2/298 amino acids are different) indicating that the coding region of *Machi3-1* is well conserved in different apple cultivars (Fig. S4). However, while the promoter regions show strong overall similarity, significant differences were found in the middle region of the promoter (Fig. 3C and Fig. S5-7). The ‘Freedom’ contains a 38 nt long insertion and a 15 nt long deletion relative to the ‘Jonagold’ promoter (Fig. 3C). More interestingly, both promoters contain 59-64 nt long direct repeat segments (referred to as boxes). However, the ‘Jonagold’ promoter contains only 2 boxes, while 5 boxes are present in the promoter of the ‘Freedom’ *Machi3-1* gene (Fig. 3C). Thus the ‘Freedom’ *Machi3-1* promoter is longer than the Jonagold promoter (1417 nt and 1202 nt, respectively). The box1 and box5 of the ‘Freedom’ promoter resemble to the box1 and box2 of the ‘Jonagold’ promoter respectively, while the ‘Freedom’ box2, 3 and 4 are more similar to each other (Fig. S6). Further studies revealed that *Machi3-1* was present in heterozygous form in both ‘Freedom’ and ‘Jonagold’ cultivars (Fig. S8), the second allele in both cultivars was a putative pseudogene. The promoter of the pseudogene (ps promoter for <u>ps</u>eudogene promoter) contained 2 boxes. The three alleles will be referred to as *5B-Machi3-1, 2B-Machi3-1* and *ps-Machi3-1*, respectively (Fig. 3C and Fig. S8).

### 5B-Machi3-1 allele co-segregate with high Machi3-1 protein level in the nectar of hybrid apples

Multiplication of a repeat region in a promoter can dramatically increase transcriptional activity (Espley *et al*., 2009). We assumed that 5 box containing *5B-Machi3-1* promoter is responsible for the strong nectary-, and stigma-specific expression of *Machi3-1* mRNA and indirectly for the nectar- and stigma-specific accumulation of Machi3-1 protein in the ‘Freedom’ cultivar. To confirm that the *5B-Machi3-1* promoter is essential for the specific expression, the F1 hybrids from ‘Freedom’ × ‘Red Rome’ and ‘Freedom’ × ‘Red Winter’ (Free. × R.R. and Free. × R.W.) crosses were studied. The Machi3-1 protein is not detectable in the nectars of the ‘Red Rome’ and ‘Red Winter’ cultivars and both are homozygous for the ps-Machi3-1 alleles (Fig. 4 and Fig. S9). The ‘Freedom’ harbors one *5B-Machi3-1* and one *ps-Machi3-1* allele and contains Machi3-1 protein in the nectar. The F1 progenies segregated close to the 1:1 for *5B-Machi3-1/ps-Machi3-1* heterozygous and for *ps-Machi3-1/ps-Machi3-1* homozygous plants (Free. × R.R. F1 segregated for 7:7, while Free. × R.W. F1 hybrids segregated for 11:13). Only 6 Free. × R. R. and 8 Free. × R. W. F1 plants developed flower in the year of the study. We found that Machi3-1 protein accumulated to easily detectable levels in the nectar of all F1 progenies that inherited the Freedom *5B-Machi3-1* allele (Fig. 4 and Fig. S9), while the progenies that inherited the ‘Freedom’ *ps-Machi3-1* allele did not accumulate the protein in their nectar (stigma samples were not collected). These results indicate that the *Machi3-1* gene is present in a single copy in ‘Freedom’, and that the *5B-Machi3-1* allele is required and sufficient for the intense nectar-specific (and likely stigma-specific) protein expression.

**Fig 4.**
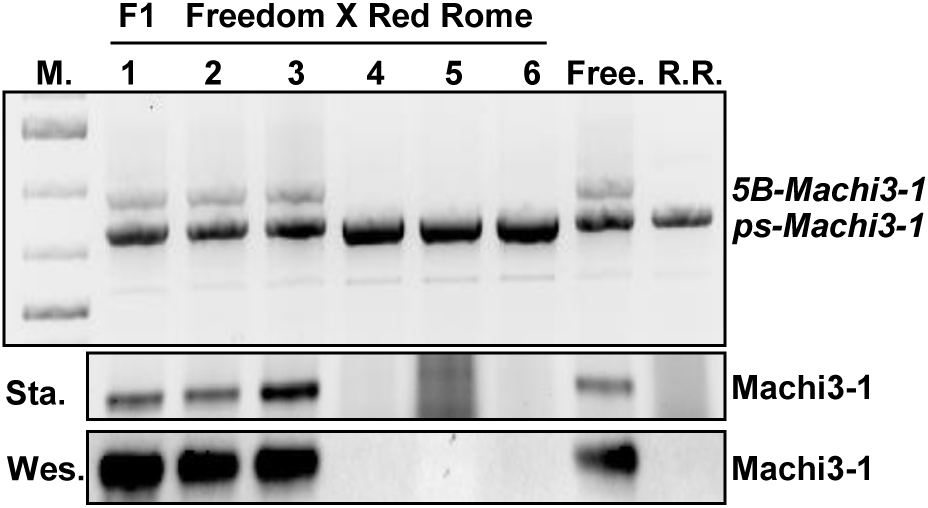
Machi3-1 nectar expression co-segregate with the *5B-Machi3-1* ‘Freedom’ allele. F1 progenies from ‘Freedom’ X ‘Red Rome’ crossing were genotyped for the *Machi3-1* alleles (upper panel). ‘Freedom’ and ‘Red Rome’ genotypes (Free. and R.R., respectively) are also shown. M. indicates DNA size marker (1.5, 1.0, 0.7 and 0.5 kb bands are shown). The accumulation of the Machi3-1 protein in the nectars of the flowering F1 plants (others did not flower) was studied by stain-free gel visualization (Sta.) and by western-blot assay (Wes.).

### The 5B-Machi3-1 promoter can confer nectary- and stigma-specific expression in tobacco

If the *trans* factors that are responsible for the nectar- and stigma-specific expression of the *5B-Machi3-1* allele in apple are also present in other dicot plants, the *5B-Machi3-1* promoter can be used as an efficient biotechnology tool for nectary- and stigma-specific expression. To test this assumption transgenic tobacco lines were generated with constructs containing the promoter and the coding region of the *5B-Machi3-1* or the *2B-Machi3-1* alleles (Fig. 5A). We found that the Machi3-1 protein was easily detectable in the nectar of 4 out of 16 *5B-Machi3-1* transgenic tobacco lines (Fig. 5A). By contrast, the transgenic protein could not be detected in any of the *2B-Machi3*-1 transgenic tobacco nectars (0/17 plants). We have also studied the accumulation of the Machi3-1 protein in the stigma of two *5B-Machi3-1* plants that expressed the Machi3-1 protein in their nectars and in two *2B-Machi3-1* plants, which did not accumulate the protein (Fig. 5C and 5D). We found that the expression in the nectar and the stigma correlated. In *5B-Machi3*-1 transgenic plants, the Machi3-1 protein accumulated to high levels both in the nectar and the stigma. By contrast, in the *2B-Machi3-1* transgenic plants the Machi3-1 protein could not be detected in either the nectar or the stigma samples. These results show that all *trans* factors that are required for the specific expression are also present in tobacco. Thus, the *5B-Machi3-1* promoter can be used for efficient nectary- and stigma-specific expression in various dicot plants.

**Fig 5.**
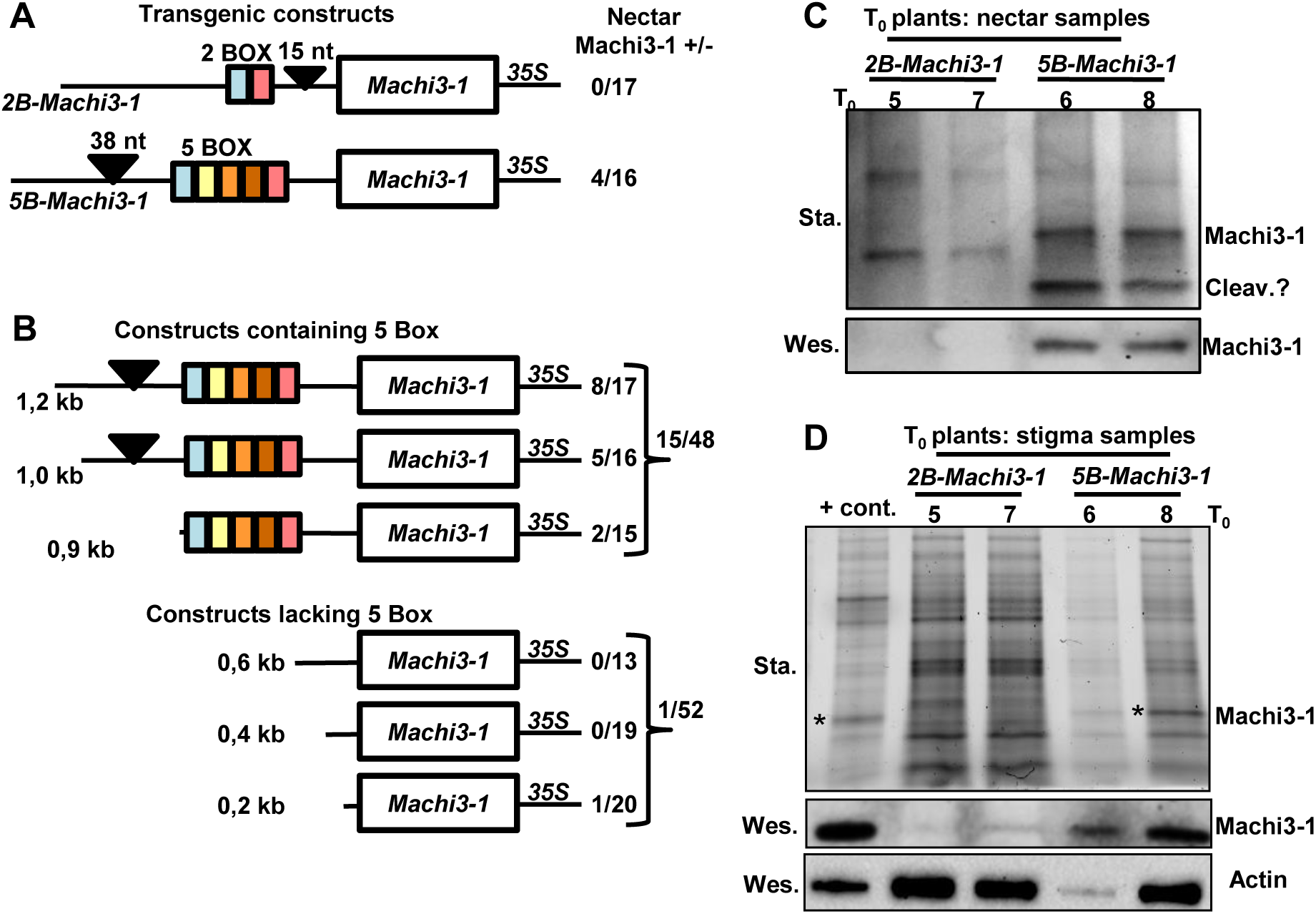
*5B-Machi3-1* transgenic tobacco plants express the Machi3-1 protein in the nectar and the stigma. (A) Non-proportional schematic representation of the *5B-Machi3-1* and *2B-Machi3-1* transgenic constructs. The promoter and the coding regions from the *5B-Machi3-1* and *2B-Machi3-1* alleles with the *35S* terminator segment were used to generate transgenic tobacco lines. The ratio of transgenic plants expressing/non-expressing Machi3-1 protein to detectable levels in the nectar is shown at the right side (nectar Machi3-1 +/-). (B) The 5 box promoter region is required for efficient expression in the nectar. Transgenic tobacco plants were generated with *5B-Machi3-1* promoter deletion constructs, and then the Machi3-1 protein expression in the nectar was tested. (C-D) Machi3-1 protein accumulates in the nectar and the stigma tissues of certain *5B-Machi3-1* transgenic tobacco plants. The accumulation of the Machi3-1 protein was studied by Stain-free gel visualization (Sta.) and by western-blot assay (Wes.). (C) The nectar profile of 2-2 independent T_0_ *2B-Machi3-1* (T_0_ plants 5 and 7) and *5B-Machi3-1* (T_0_ plants 6 and 8) transgenic plants. We selected *5B-Machi3-1* plants, which accumulate the transgenic protein in the nectar. Cleav. shows putative cleavage fragment of the Machi3-1 protein. (D) The protein profile of stigma samples of the same transgenic lines. ‘Freedom’ apple stigma extract was run as positive control (+cont.). * marks Machi3-1 band in the stigma samples. Note that Machi3-1 protein is one of the most abundant protein in ‘Freedom’ stigma sample. Actin probe was used as loading control for the western-blot. Note that although stigma sample of *5B-Machi3-1* plant 6 is underloaded, the Machi3-1 protein is still easily detectable.

### The 5 box region of the Machi3-1 promoter is required for efficient expression

To directly prove that the boxes of the *5B-Machi3-1* promoter are required for the specific expression, plants were transformed with deletion constructs generated from the *5B-Machi3-1* plasmid (Fig. 5B), and then we studied the accumulation of the Machi3-1 protein in the nectars of the transformants (stigma samples were not collected in this experiment). The results suggest that the 5 boxes are essential for the efficient expression, only 1 out of 52 plants accumulated Machi3-1 protein in the nectar at detectable levels when constructs lacking the 5 boxes were used (we combined the results of 0.6, 0.4 and 0.2 constructs, see Fig. 5B). By contrast, 15/48 plants expressed Machi3-1 protein in the nectar (Fig. 5B) when the promoter contained the 5 box region (combining the results of 1.2, 1.0 and 0.9 constructs).

### MYB305 could play an important role in the regulation of 5B-Machi3-1

As the 5 box promoter region is required for the efficient and specific expression of the *5B-Machi3-1* gene, we assumed that transcription factors that selectively bind to this region play important role in the regulation. To identify 5 box region specific transcription factor binding sites, we compared the 5 box and 2 box promoter regions of the *5B-Machi3-1* and *2B-Machi3-1* alleles. Interestingly, we identified 4 potential MYB binding sites in the 5 box and only 1 in the 2 box promoter region (Fig. S7). In tobacco, the MYB305 transcription factor directs the nectary-specific expression of many genes including nectarins (Liu *et al*., 2009). We hypothesized that MYB305 and the apple homolog of MYB305 (referred to as MdMYB305, gene: MDP0000344978) directs the expression of 5B-Machi3-1 in the nectary as well as in the stigma cells. If these assumptions are correct, MYB305 and *5B-Machi3-1* (but not *2B-Machi3-1*) mRNAs are co-expressed. To test it, qRT-PCR and semi qRT-PCR assays were conducted to monitor the expression of *MdMYB305* in ‘Freedom’ and ‘Jonagold’ cultivars (Fig. 6A and Fig. S10A). The *MdMYB305* mRNA expressed similarly in both cultivars, it was abundant in the nectary and stigma samples but was barely detectable in leaves (Fig. 6A and Fig. S10A). As the ‘Freedom’ *5B-Machi3-1* (but not the ‘Jonagold’ *2B-Machi3-1*) mRNAs expressed similarly (Fig. 3B and Fig. S10A), we concluded that in apple the *MdMYB305* transcript is co-expressed with the *5B-Machi3-1* but not with the *2B-Machi3-1* mRNA. In line with our results, a recent RNA-seq experiment showed that in ‘Golden Delicious’ cultivar (SRA-NCBI: SRP125281 study), which contains one copy of each of the *2B-Machi3-1* and the *ps-Machi3-1* alleles, the *MdMYB305* mRNA expressed to high levels in the stigma and style and that, *2B-Machi3-1* transcript accumulated to low levels in these tissues (Fig. S10B.).

**Fig 6.**
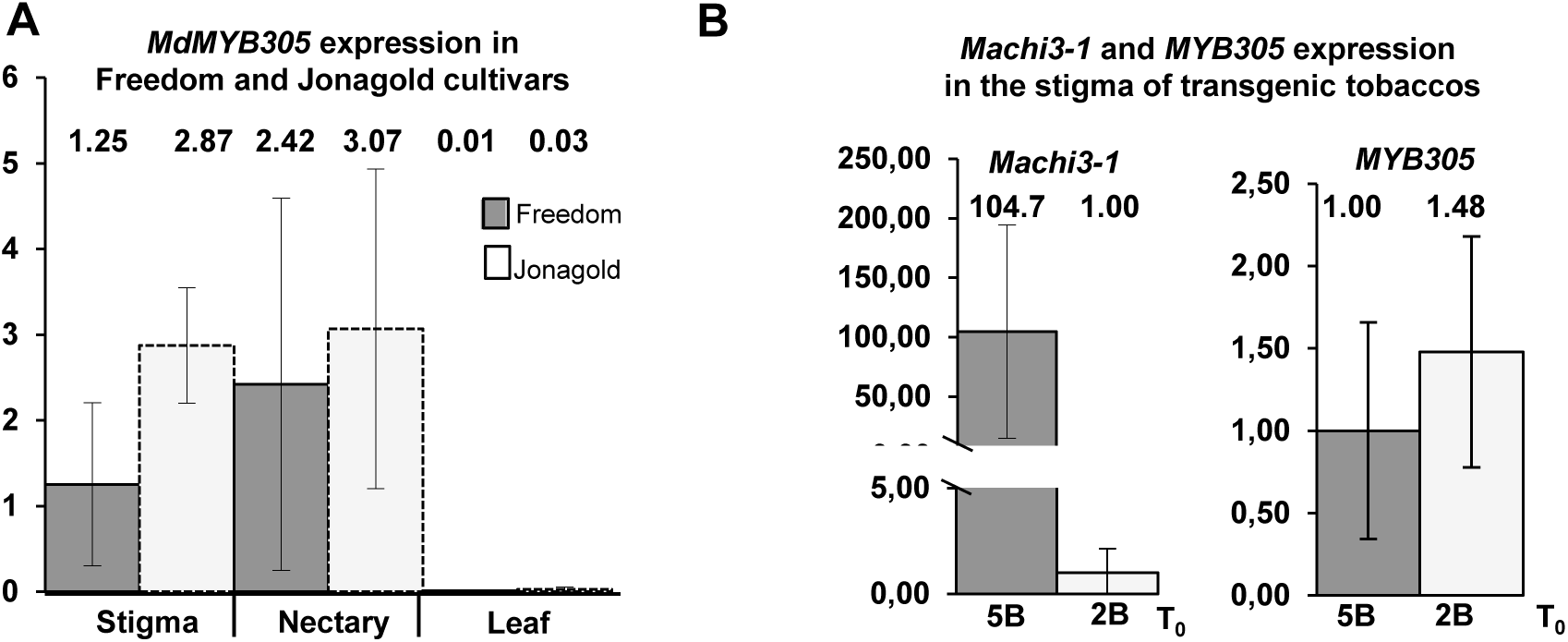
*5B-Machi3-1* is co-expressed with *MYB305*. (A) In apple, the *MdMYB305* is expressed in the nectary and the stigma but not in the leaf. qRT-PCR was conducted to monitor the expression of *MdMYB305* in the nectary, stigma and leaf samples of ‘Freedom’ and ‘Jonagold’ cultivars. Note that *MdMYB305* expressed to comparable level in both cultivars. (B) In *5B-Machi3-1* transgenic tobacco, *Machi3-1* mRNA is co-expressed with *MYB305* transcript in the stigma. qRT-PCR analysis of the expression of *MYB305* and *Machi3-1* mRNAs in the stigma of *5B-Machi3-1* (5B) and *2B-Machi3-1* (2B) transgenic tobaccos.

To further support that *5B-Machi3-1* and *MYB305* transcripts are co-expressed, *Machi3-1* and *MYB305* mRNA levels were studied in the stigma tissues of *5B-Machi3-1* and *2B-Machi3-1* transgenic tobaccos (Fig. 6B and Fig. S10C). The *MYB305* mRNA expressed to high levels in the stigma of both transgenic lines. Relevantly, the *5B-Machi3-1* transgenic mRNA was also abundant in the stigma, whereas the *2B-Machi3-1* mRNA expressed to low levels (Fig. 6B and Fig. S10C). The fact that the *MdMYB305* and *MYB305* mRNAs co-express only with the *5B-Machi3-1* transcript, and that it is transcribed from the 4 MYB binding site containing *5B-Machi3-1* gene suggests that MYB305 homologs play an important role in the regulation of *5B-Machi3-1* expression.

### The 5B-Machi3-1 might be introgressed from the Malus floribunda 821 to different cultivars

We found that the ‘Freedom’ cultivar contains the *5B-Machi3-1* allele in heterozygous form, while ‘Red Rome’, ‘Red Winter’ and ‘Jonagold’ cultivars harbored the *2B-Machi3-1* and/or the *ps-Machi3-1* alleles. To clarify how widespread is the *5B-Machi3-1* allele, we PCR genotyped several more apple cultivars (Fig. 7A and Fig. S11). We found that the *5B-Machi3-1* allele is present in heterozygous or homozygous form in the genome of many (but not all) apple cultivars that contain the *M.floribunda* 821 derived *Vf* scab resistance gene, while it was not found in the cultivars that did not harbor the *Vf* gene (Gessler and Pertot, 2012). The Machi3-1 protein was present in the nectar of all the *5B-Machi3-1* allele containing cultivars (Fig. 7A) but it was not detectable in the nectar of cultivars lacking the *5B-Machi3-1* allele. These data confirm that the *5B-Machi3-1* allele is sufficient for the accumulation of Machi3-1 protein in the nectar.

**Fig 7.**
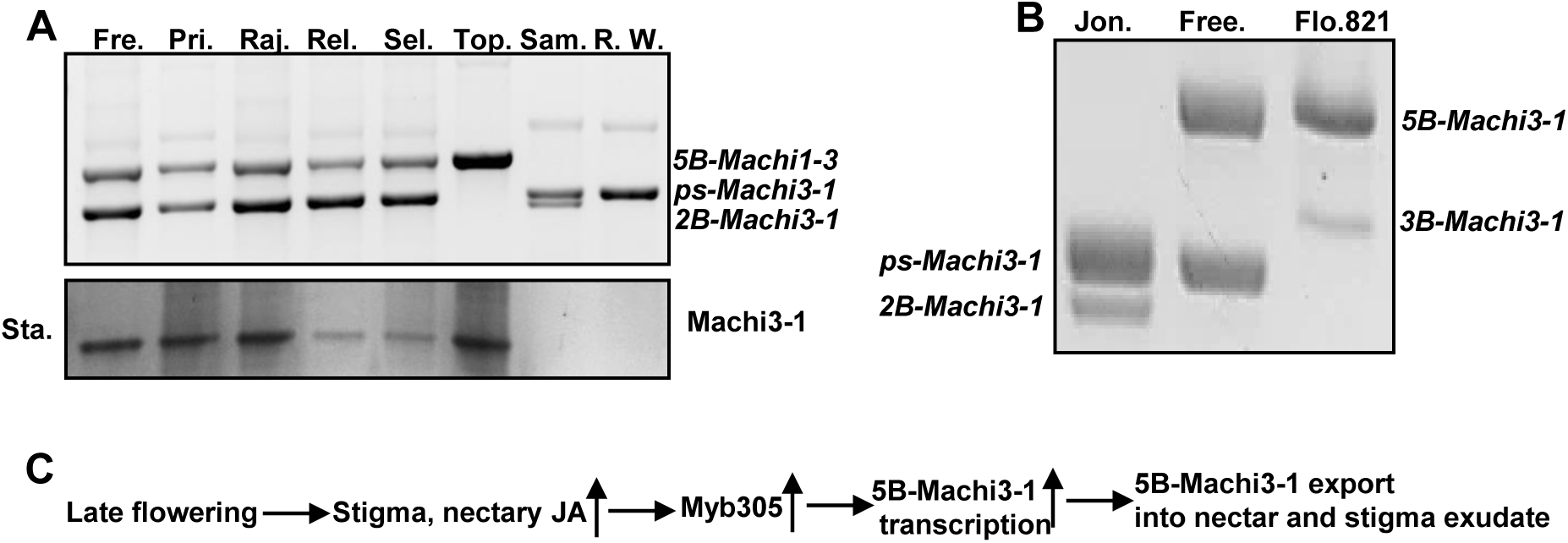
The *5B-Machi3-1* allele containing apple cultivars express the Machi3-1 protein in the nectar. (A) The Machi3-1 protein is abundant in the nectar of *5B-Machi3-1* allele containing apple cultivars Different apple cultivars were genotyped for *Machi3-1* and their nectar samples were stain-free visualized (Sta.) (Fre.-‘Freedom’, Pri.-‘Prima’, Raj.- ‘Rajka’, Rel.-’Releika’, Sel.-‘Selena’, Top.-‘Topaz’, Sam.-‘Sampion’, R.W.-‘Red Winter’). (B) *M.floribunda* 821 contains a *5B-Machi3-1* allele. (C) Model of regulation of *Machi3-1* expression.

Various Vf scab resistant apple cultivars including ‘Freedom’, ‘Releika’ and ‘Topaz’ contain the *5B-Machi3-1* allele (Fig. S11). Although these cultivars were generated in different breeding programs in the USA (‘Freedom’), Germany (’Releika’) and Czech Republic (‘Topaz’), progenies from the *M.floribunda* 821 clone and ‘Rome Beauty’ (F_2_26829-2-2) crossing were used in each programs (Gessler and Pertot, 2012). Therefore, we postulated that the *5B-Machi3-1* allele was introgressed from the *M.floribunda* 821 ancestor. Indeed, we found that the *M.floribunda* 821 contains a *5B-Machi3-1* allele (Fig. 7B), which is almost identical (295/298 amino acids of the predicted proteins are identical) to the Freedom *5B-Machi3-1* allele (Fig. S12). The promoter of *M.floribunda* 821 *5B-Machi3-1* is also highly similar to the promoter of the ‘Freedom’ *5B-Machi3-1* gene, it contains the 5 boxes and all 4 MYB binding sites are present (Fig. S12 and S13). We hypothesize that Machi3-1 protein is also abundant in the nectar and the stigma of *M.floribunda* 821 (flowers were not available to test this prediction). *Machi3-1* is present in heterozygous form in *M.floribunda* 821, the second *Machi3-1* allele has a specific promoter with 3 boxes (*3B-Machi3-1*) (Fig. 7B). Taken together, these data indicate that the *5B-Machi3-1* allele was introgressed from the *M.floribunda* 821 clone into various apple cultivars.

## Discussion

Here we show that the *5B-Machi3-1* allele, which was introgressed from the *M.floribunda* 821 ancestor into certain apple cultivars, encodes an acidic chitinase III protein that accumulates to very high levels in the stigma and the nectar, the primary niches of the *Erwinia* infection. As its protein product inhibits *in vitro* the growth and biofilm formation of *Erwinia* at physiological concentration, we hypothesize that the Machi3-1 protein could partially protect apple against *Erwinia* infection and defend nutritious flower secretions from microbial infections.

### Regulation of the expression of the 5B-Machi3-1 allele

We found that the *Machi3-1* acidic chitinase III gene is present in at least three different forms in various apple cultivars, one is a pseudogene, while the two other alleles (*5B-Machi3-1* and *2B-Machi3-1*) encode very similar proteins (Fig. S4). However, the regulation of the two active alleles is markedly different. The *2B-Machi3-1* allele expresses to low levels in all studied tissues, while the *5B-Machi3-1* mRNAs shows very intense expression in the nectary and stigma (Fig. 3). Our segregation, association and transgenic assays demonstrate that the 5 boxes of the *5B-Machi3-1* promoter is required for the nectary- and stigma-specific transcript and for the nectar- and stigma-specific protein expressions (Fig. 4, 5 and 7). Previously it was shown that MYB305 (or its homologs as MYB21 and MYB24) transcription factor plays a critical role in the expression of nectar proteins in tobacco, snapdragon and Arabidopsis (Roy *et al*., 2017). MYB305 is activated by JA (and probably by auxin), then it binds to the promoters of many nectary-specific genes (including certain nectarins) and promotes their transcription (MYB305 might also stimulate indirectly the accumulation of other nectar proteins). We identified four potential MYB binding sites on the 5 box region of the 5B-Machi3-1 allele but only one on the 2B promoter (Fig. S7). We demonstrated that in addition to the nectary (Liu *et al*., 2009), tobacco *MYB305* is also expressed to high levels in the stigma (Fig. 6). Moreover, we showed that the apple homolog *MdMYB305* transcripts are also abundant in the nectary and stigma tissues, while it accumulates to low levels in the leaf (Fig. 6). We propose that the *5B-Machi3-1* gene is similarly regulated in apple and transgenic tobacco plants (Fig. 7C). During late phase of flower development, JA increases the MYB305 level in the nectary and probably in the stigma, and then MYB305 binds to the 5 box region and promotes the transcription of *5B-Machi3-1* gene. As Machi3-1 protein contains a signal peptide, it can be secreted into the nectar explaining why *5B-Machi3-1* mRNA is abundant in the nectary, while the Machi3-1 protein accumulates in the nectar. We hypothesize that Machi3-1 protein is also secreted from the stigma cells. As both apple and tobacco have wet stigma (Sang *et al*., 2012; Losada and Herrero, 2012), we assume that in both plants Machi3-1 protein is secreted into the stigma exudate (Fig. 7C) (also see below).

Interestingly, the strong expression of *MYB305* in nectary and stigma is not restricted to the plants having wet stigma such as tobacco or apple. The Arabidopsis homolog MYB21 is also expressed to high levels in both nectary and in the papilla cells of the dry stigma (Osaka *et al*., 2013).

### Machi3-1 protein might interfere with Erwinia infection at two different steps

We found that the Machi3-1 acidic chitinase III protein is present at very high concentration in the nectar and the stigma of the *5B-Machi3-1* allele containing cultivars (Fig. 3 and 7). During apple infection, *Erwinia* propagates first in the stigma exudate, then in the nectar and finally enters into the plant through the stomata of the nectary (Bubán *et al*., 2003; Farkas *et al*., 2012). As biofilm formation is critical for the fruit and shoot infection of *Erwinia* (Koczan *et al*., 2011), it is likely that biofilm formation is also required for efficient flower infection. *In vitro*, the Machi3-1 protein inhibits growth and biofilm formation of *Erwinia* at a concentration it is present in the nectar of *5B-Machi3-1* allele containing apple cultivars (Fig. 2). These findings suggest that the Machi3-1 protein can interfere with the propagation and infection of *Erwinia* in the nectar of the *5B-Machi3-1* allele containing cultivars. Moreover, the Machi3-1 protein might also inhibit *Erwinia* infection at the stigma. Machi3-1 was one of the most abundant protein in the ‘Freedom’ stigma sample (Fig. 5D, see the + control sample and Fig. S3) that contains both the stigma tissue and exudate. Machi3-1 has a signal peptide, therefore we assume that it is secreted from the stigma cells and accumulates in the exudate. The Machi3-1 protein might be so abundant in the stigma exudate that it can interfere with the propagation of *Erwinia*. Thus we propose that accumulation of Machi3-1 protein could protect apples from *Erwinia* infection by forming two barriers, it interferes with the replication and biofilm formation of the *Erwinia* in the stigma exudate as well as in the nectar, thereby protecting the plants. Our data shows that the *5B-Machi3-1* allele was introduced into different cultivars from *M.floribunda* 821 clone. We propose that the inhibitory Machi3-1 protein level in the stigma exudate and the nectar could contribute to the partial *Erwinia* resistance of these *M.floribunda* 821 derived cultivars. Importantly, if *5B-Machi3-1* contributes to the *Erwinia* resistance, this effect is not detectable in the frequently used shoot inoculation assays. Finally, as chitinases have wide spectrum antimicrobial effect (Cletus *et al*., 2013), the high concentration of Machi3-1 protein could effectively protect the nectar and the stigma exudate from various microbial infection. Thus it can keep the optimal composition of these important fluid secretions, thereby enhancing the efficiency of pollination and fertilization.

## Supplementary Data

**Supplementary Table S1.** List of primers.

**Supplementary Materials and methods S1.** Protein sequencing.

**Fig. S1.** Homology modelling of Machi3-1 and Hevamine proteins by Swiss-PdbViewer.

**Fig. S2.** Machi3-1 has a very weak lysozyme activity.

**Fig. S3.** Machi3-1 expresses to high levels in the stigma and the nectar of ‘Freedom’ apple cultivar.

**Fig. S4.** Alignment of the coding regions of *5B-Machi3-1* and *2B-Machi3-1* genes.

**Fig. S5.** Similarity of the promoter regions of *5B-Machi3-1* and *2B-Machi3-1* genes.

**Fig. S6.** Comparison of the boxes from the *5B-Machi3-1* and *2B-Machi3-1* promoters.

**Fig. S7.** Predicted transcription factor binding sites in the 5 box and 2 box regions.

**Fig. S8.** The structure of the three *Machi3-1* alleles.

**Fig. S9.** Machi3-1 protein expression co-segregate with the *5B-Machi3-1* ‘Freedom’ allele.

**Fig. S10.** The *MYB305* and *5B-Machi3-1* transcripts are co-expressed.

**Fig. S11.** Genotyping of different apple cultivars for *Machi3-1* alleles.

**Fig. S12.** The **‘**Freedom’ and *M.floribunda* 821 *5B-Machi3-1* genes are highly similar.

**Fig. S13.** Predicted transcription factor binding sites in the 5 box region of the promoter of *M.floribunda* 821 *5B-Machi3-1* gene.

## Abbreviations

Machi3-1: *Malus* acidic chitinase 3-1 protein

## Acknowledgements

We are grateful to M. Peline Toth, M. Csanyi and J. Nadudvarine Novak for transformations and various technical assistance. We thanks A. Kerekes (Agricultural Biotechnology Institute) for the anti-Machi3-1 antibody and A. Auber (Agricultural Biotechnology Institute) for his help with the bioinformatical studies and Z. K. Varga for his help in statistical analyses. We are especially grateful to E. Van de Weg (Wageningen, Netherlands) for the *M.floribunda* 821 DNA samples. We thank A. Hegedus (Szent Istvan University, Hungary) for the useful comments about the manuscript. Research was supported by grants from the OTKA (K81481, CK80029). A. Kurilla is a graduate student of ELTE “Classical and Molecular Genetics” PhD program.

## References

Adrangi S, Faramarzi MA. 2013. From bacteria to human: A journey into the world of chitinases. Biotechnology Advances 31, 1786–1785.

Bender RL, Fekete ML, Klinkenberg PM, et al. 2013. PIN6 is required for nectary auxin response and short stamen development. The Plant JournalL 74, 893–904.

Bevan MW, Mason SE, Goelet P. 1985. Expression of tobacco mosaic virus coat protein by a cauliflower mosaic virus promoter in plants transformed by Agrobacterium. The EMBO Journal 4, 1921–6.

Bowman JL, Smyth DR. 1999. CRABS CLAW, a gene that regulates carpel and nectary development in Arabidopsis, encodes a novel protein with zinc finger and helix-loop-helix domains. Development (Cambridge, England) 126, 2387–96.

Bubán T, Orosz-Kovács Z, Farkas Á. 2003. The nectary as the primary site of infection by Erwinia amylovora (Burr.) Winslow et al.: A mini review. Plant Systematics and Evolution 238, 183–194.

Carter C, Healy R, O’Tool NM, Naqvi SMS, Ren G, Park S, Beattie G a, Horner HT, Thornburg RW. 2007. Tobacco nectaries express a novel NADPH oxidase implicated in the defense of floral reproductive tissues against microorganisms. Plant physiology 143, 389–399.

Chung M-C, Dean S, Marakasova ES, Nwabueze AO, van Hoek ML. 2014. Chitinases Are Negative Regulators of Francisella novicida Biofilms (MF Feldman, Ed.). PLoS ONE 9, e93119.

Chung KP, Zeng Y. 2017. An Overview of Protein Secretion in Plant Cells. Methods in molecular biology (Clifton, N.J.). 19–32.

Cletus J, Balasubramanian V, Vashisht D, Sakthivel N. 2013. Transgenic expression of plant chitinases to enhance disease resistance. Biotechnology Letters 35, 1719–1732.

Edlund AF, Swanson R, Preuss D. 2004. Pollen and stigma structure and function: the role of diversity in pollination. The Plant cell 16 Suppl, S84–97.

Espley R V., Brendolise C, Chagne D, et al. 2009. Multiple Repeats of a Promoter Segment Causes Transcription Factor Autoregulation in Red Apples. the Plant Cell Online 21, 168–183.

Farkas Á, Mihalik E, Dorgai L, Bubán T. 2012. Floral traits affecting fire blight infection and management. Trees - Structure and Function 26, 1–20.

Ferrari AR, Gaber Y, Fraaije MW. 2014. A fast, sensitive and easy colorimetric assay for chitinase and cellulase activity detection. Biotechnology for biofuels 7, 37.

Gessler C, Pertot I. 2012. Vf scab resistance of Malus. Trees - Structure and Function 26, 1–14.

Gusberti M, Klemm U, Meier MS, Maurhofer M, Hunger-Glaser I. 2015. Fire blight control: The struggle goes on. a comparison of different fire blight control methods in switzerland with respect to biosafety, efficacy and durability. International Journal of Environmental Research and Public Health 12, 11422–11447.

Heil M. 2011. Nectar: Generation, regulation and ecological functions. Trends in Plant Science 16, 191–200.

Jin J, Tian F, Yang D-C, Meng Y-Q, Kong L, Luo J, Gao G. 2017. PlantTFDB 4.0: toward a central hub for transcription factors and regulatory interactions in plants. Nucleic acids research 45, D1040–D1045.

Kelley DR, Estelle M. 2012. Ubiquitin-mediated control of plant hormone signaling. Plant Physiology 160, 47–55.

Koczan JM, Lenneman BR, McGrath MJ, Sundin GW. 2011. Cell surface attachment structures contribute to biofilm formation and xylem colonization by Erwinia amylovora. Applied and Environmental Microbiology 77, 7031–7039.

Koczan JM, McGrath MJ, Zhao Y, Sundin GW. 2009. Contribution of Erwinia amylovora exopolysaccharides amylovoran and levan to biofilm formation: implications in pathogenicity. Phytopathology 99, 1237–44.

De la Barrera E, Nobel PS. 2004. Nectar: properties, floral aspects, and speculations on origin. Trends in Plant Science 9, 65–9.

Lee J-Y, Baum SF, Oh S-H, Jiang C-Z, Chen J-C, Bowman JL. 2005. Recruitment of CRABS CLAW to promote nectary development within the eudicot clade. Development (Cambridge, England) 132, 5021–32.

Liu G, Ren G, Guirgis A, Thornburg RW. 2009. The MYB305 Transcription Factor Regulates Expression of Nectarin Genes in the Ornamental Tobacco Floral Nectary. The Plant Cell Online 21, 2672–2687.

Liu G, Thornburg RW. 2012. Knockdown of MYB305 disrupts nectary starch metabolism and floral nectar production. The Plant JournalL 70, 377–88.

Losada JM, Herrero M. 2012. Arabinogalactan-protein secretion is associated with the acquisition of stigmatic receptivity in the apple flower. Annals of Botany 110, 573–584.

Lv C, Masuda T, Yang H, Sun L, Zhao G. 2011. High-capacity calcium-binding chitinase III from pomegranate seeds (Punica granatum Linn.) is located in amyloplasts. Plant Signaling and Behavior 6, 1963–1965.

Ma XL, Milne RI, Zhou HX, Fang JY, Zha HG. 2017. Floral nectar of the obligate outcrossing Canavalia gladiata (Jacq.) DC. (Fabaceae) contains only one predominant protein, a class III acidic chitinase. (HP Mock, Ed.). Plant Biology (Stuttgart, Germany) 19, 749–759.

Malnoy M, Martens S, Norelli JL, Barny M-A, Sundin GW, Smits THM, Duffy B. 2012. Fire Blight: Applied Genomic Insights of the Pathogen and Host. Annual Review of Phytopathology 50, 475–494.

Masuda T, Zhao G, Mikami B. 2015. Crystal structure of class III chitinase from pomegranate provides the insight into its metal storage capacity. Bioscience, Biotechnology and Biochemistry 79, 45–50.

Migh E, Götz T, Földi I, et al. 2018. Microtubule organization in presynaptic boutons relies on the formin DAAM. Development 145, dev158519.

Mihalik E, Radvánszky A, Dorgai L, Bubán T. 2004. Study of Erwinia amylovora colonization and migration on blossoms of susceptible and tolerant apple cultivars. International Journal of Horticultural Science (Hungary) 10, 15–19.

Min Y, Bunn JI, Kramer EM. 2019. Homologs of the STYLISH gene family control nectary development in Aquilegia. The New Phytologist 221, 1090–1100.

Morel P, Heijmans K, Ament K, Chopy M, Trehin C, Chambrier P, Rodrigues Bento S, Bimbo A, Vandenbussche M. 2018. The Floral C-Lineage Genes Trigger Nectary Development in Petunia and Arabidopsis. The Plant Cell 30, 2020–2037.

Nash JA, Ballard TNS, Weaver TE, Akinbi HT. 2006. The Peptidoglycan-Degrading Property of Lysozyme Is Not Required for Bactericidal Activity In Vivo. The Journal of Immunology 177, 519–526.

Nogueira FCS, Farias ARB, Teixeira FM, Domont GB, Campos FAP. 2018. Common Features Between the Proteomes of Floral and Extrafloral Nectar From the Castor Plant (Ricinus Communis) and the Proteomes of Exudates From Carnivorous Plants. Frontiers in Plant Science 9, 549.

O’Toole GA. 2011. Microtiter Dish Biofilm Formation Assay. Journal of Visualized Experiments.

Osaka M, Matsuda T, Sakazono S, et al. 2013. Cell Type-Specific Transcriptome of Brassicaceae Stigmatic Papilla Cells From a Combination of Laser Microdissection and RNA Sequencing. Plant and Cell Physiology 54, 1894–1906.

Piqué N, Miñana-Galbis D, Merino S, Tomás JM. 2015. Virulence factors of Erwinia amylovora: A review. International Journal of Molecular Sciences 16, 12836–12854.

Pusey PL, Rudell DR, Curry EA, Mattheis JP. 2008. Characterization of stigma exudates in aqueous extracts from apple and pear flowers. HortScience 43, 1471–1478.

Radhika V, Kost C, Boland W, Heil M. 2010. The role of jasmonates in floral nectar secretion. (A Rahman, Ed.). PloS One 5, e9265.

Rejón JD, Delalande F, Schaeffer-Reiss C, Carapito C, Zienkiewicz K, Alché J de D, Rodríguez-García MI, Van Dorsselaer A, Castro AJ. 2014. The plant stigma exudate. A biochemically active extracellular environment for pollen germination? Plant Signaling & Behavior 9, e28274.

Rejón JD, Delalande F, Schaeffer-Reiss C, Carapito C, Zienkiewicz K, de Dios Alché J, Rodríguez-García MI, Van Dorsselaer A, Castro AJ. 2013. Proteomics profiling reveals novel proteins and functions of the plant stigma exudate. Journal of Experimental Botany 64, 5695–5705.

Roy R, Schmitt AJ, Thomas JB, Carter CJ. 2017. Review: Nectar biology: From molecules to ecosystems. Plant Science 262, 148–164.

Sang YL, Xu M, Ma FF, Chen H, Xu XH, Gao X-Q, Zhang XS. 2012. Comparative proteomic analysis reveals similar and distinct features of proteins in dry and wet stigmas. PROTEOMICS 12, 1983–1998.

Sasu MA, Wall KL, Stephenson AG. 2010. Antimicrobial nectar inhibits a florally transmitted pathogen of a wild Cucurbita pepo (Cucurbitaceae). American Journal of Botany 97, 1025–1030.

Schmitt AJ, Roy R, Klinkenberg PM, Jia M, Carter CJ. 2018a. The Octadecanoid Pathway, but Not COI1, Is Required for Nectar Secretion in Arabidopsis thaliana. Frontiers in Plant Science 9.

Schmitt AJ, Sathoff AE, Holl C, Bauer B, Samac DA, Carter CJ. 2018b. The major nectar protein of Brassica rapa is a non-specific lipid transfer protein, BrLTP2.1, with strong antifungal activity. Journal of Experimental Botany 69, 5587–5597.

Shen C-R, Chen Y-S, Yang C-J, Chen J-K, Liu C-L. 2010. Colloid Chitin Azure Is a Dispersible, Low-Cost Substrate for Chitinase Measurements in a Sensitive, Fast, Reproducible Assay. Journal of Biomolecular Screening 15, 213–217.

Silhavy D, Molnár A, Lucioli A, Szittya G, Hornyik C, Tavazza M, Burgyán J. 2002. A viral protein suppresses RNA silencing and binds silencing-generated, 21-to 25-nucleotide double-stranded RNAs. The EMBO Journal 21, 3070–80.

Stitz M, Hartl M, Baldwin IT, Gaquerel E. 2014. Jasmonoyl-L-isoleucine coordinates metabolic networks required for anthesis and floral attractant emission in wild tobacco (Nicotiana attenuata). The Plant Cell 26, 3964–83.

Szittya G, Molnár A, Silhavy D, Hornyik C, Burgyán J. 2002. Short defective interfering RNAs of tombusviruses are not targeted but trigger post-transcriptional gene silencing against their helper virus. The Plant Cell 14, 359–72.

Tanveer T, Shaheen K, Parveen S, Kazi AG, Ahmad P. 2014. Plant secretomics: identification, isolation, and biological significance under environmental stress. Plant Signaling & Behavior 9, e29426.

Terwisscha Van Scheltinga AC, Hennig M, Dijkstra BW. 1996. The 1.8 Å resolution structure of hevamine, a plant chitinase/lysozyme, and analysis of the conserved sequence and structure motifs of glycosyl hydrolase family 18. Journal of Molecular Biology 262, 243–257.

Zha H-GG, Milne RI, Zhou H-XX, Chen X-YY, Sun H. 2016. Identification and cloning of class II and III chitinases from alkaline floral nectar of Rhododendron irroratum, Ericaceae. Planta 244, 805–818.

